# Riemannian Tensor Analysis Identifies a Bifurcation State in the Single-Cell Transcriptomic Landscape of Parkinson’s Disease

**DOI:** 10.64898/2026.05.09.724002

**Authors:** Moonseok Choi, Jino Choi, Kyusung Kim, Sarah Bauermeister, Do-Geun Kim

## Abstract

The transition from healthy aging to Parkinson’s disease (PD) involves highly volatile transcriptomic rewiring that remains invisible to conventional mean-based analyses. To decode this critical tipping point, we integrated single-cell RNA-sequencing with a novel Log-Euclidean Riemannian tensor framework. By conceptualizing distinct transcriptomic states as symmetric positive definite (SPD) covariance tensors, we bypassed Euclidean geometric artifacts to accurately map the macroscopic network architecture of the human prefrontal cortex. Our analysis identified a highly unstable, intermediate bifurcation (BIF) state. Thermodynamic and topological validation demonstrated that this BIF state operates as a definitive geometric saddle point, characterized by maximal von Neumann entropy and positioned perfectly equidistant between the healthy (HC) and pathological (PD) Fréchet means on the non-Euclidean manifold. Furthermore, spectral deconstruction of differential covariance networks 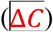 isolated the exact topological drivers of this transition—revealing the catastrophic structural collapse of core synaptic and electrophysiological hubs prior to overt pathological commitment. Ultimately, this *ab initio* geometric framework fundamentally redefines PD progression, providing a quantitative blueprint to intercept neurodegenerative trajectories during their final reversible window.

## Main

Neurodegenerative disorders, such as Parkinson’s disease (PD), are characterized by insidious pathological cascades that often precede overt clinical symptoms by decades ^1^. Long before irreversible neuronal apoptosis occurs, cells undergo subtle structural rewiring and synaptic deterioration ^2^. Consequently, identifying the precise pre-disease boundary—the critical “tipping point” that separates healthy aging from a committed pathological trajectory—remains the paramount goal in modern neurobiology ^3^. However, defining this transitional boundary has proven elusive due to the lack of analytical frameworks capable of capturing the macroscopic volatility of cellular systems before gross transcriptional collapse.

While high-dimensional single-cell transcriptomic (scRNA-seq) data encode not only mean expression levels but also the higher-order structure of gene-gene relationships, most current analyses rely on scalar representations ^4, 5^. Conventional approaches—such as differential expression testing, Euclidean dimensionality reduction, and discrete clustering—inherently reduce this biological complexity to average expression levels and static cell states ^6, 7^. While effective for identifying dominant pathological signatures, these mean-based frameworks are fundamentally blind to transitional or unstable states that manifest primarily as changes in variability, covariance, or bimodal distributional structures ^8-11^. For example, two distinct cellular states may exhibit nearly identical mean expression profiles while differing substantially in variance—a subtle discrepancy that often reflects underlying regulatory instability but remains mathematically penalized and invisible to standard comparative methods.

These limitations become especially critical when studying disease progression, where early transitional states may not involve massive transcriptional shifts, but rather emerge as subtle disruptions in gene network coordination ^12^. From a dynamical systems perspective, stable cellular states correspond to attractors in a regulatory energy landscape. Transitions between these attractors must proceed through regions of profound instability, known as bifurcation points ^13, 14^. Such critical transitions are inherently characterized by increased variance, bimodal expression, and critical slowing down—collectively recognized as early warning signals of impeding state change ^10, 15, 16^. Existing computational approaches, including trajectory inference, are primarily designed to capture smooth, directional transitions in mean expression space, rendering these highly volatile, instability-driven intermediate phases effectively undetectable ^17, 18^, and thereby creating a profound analytical blind spot that obscures the exact critical window where therapeutic intervention could potentially halt irreversible disease commitment.

Addressing this blind spot requires a fundamental conceptual shift: to accurately capture state transitions, we must decode the structure of gene-gene coordination using covariance matrices.

Because covariance matrices are symmetric positive definite (SPD) objects, they naturally form tensors that describe the geometric architecture of transcriptomic states ^19, 20^. Unlike vectors in a flat Euclidean space, SPD tensors reside on a curved Riemannian manifold. Applying conventional Euclidean operations to these tensors severely distorts their intrinsic structure and yields biologically misleading results ^21, 22^. Drawing inspiration from macroscopic neuroimaging, where diffusion tensors are robustly analyzed via Riemannian metrics ^20, 21, 23^. Applying conventional Euclidean operations to these tensors severely distorts their intrinsic structure and yields biologically misleading results. By synthesizing core mathematical principles from information geometry and statistical mechanics—specifically Log-Euclidean Fréchet mean estimation ^21^ and von Neumann entropy ^24, 25^, we engineered a fundamentally novel Riemannian tensor-based analytical framework for scRNA-seq data. By natively mapping transcriptomic states as SPD covariance tensors, our paradigm completely circumvents Euclidean distortions, enabling the precise geometric quantification of macroscopic network volatility and state transitions.

Applying this geometric framework to human prefrontal cortex scRNA-seq data from PD patients ^26^, we identify a previously unrecognized bifurcation (BIF) state positioned precisely at the tipping point between healthy aging (HC) and disease (PD). This transitional phase is characterized by massive transcriptional variance, bimodal expression of core tensor hub drivers (e.g., CNTNAP2, GRIK1, TNR), and peak network entropy—the ultimate thermodynamic hallmarks of a system on the verge of structural collapse. Strikingly, while completely invisible to conventional mean-based analyses, the BIF state emerges definitively as a high-energy ‘saddle point’ on the Riemannian manifold, located at a near-perfect equidistance between the healthy and pathological Fréchet means. Ultimately, this study establishes Riemannian tensor geometry as a computational vanguard for decoding highly volatile state transitions, providing a fundamentally new blueprint for intercepting neurodegeneration at its earliest, reversible window.

## Results

### Identification of a Critical Bifurcation State in Parkinson’s Disease via Riemannian Tensor Geometry

To characterize the transcriptomic architecture underlying the progression of PD, we first analyzed a publicly available scRNA-seq dataset derived from the human prefrontal cortex. Initial broad transcriptomic profiling using t-distributed stochastic neighbor embedding (t-SNE) visualization successfully captured the cellular landscape across HC and PD conditions (Fig. 1a). Subsequent cell-type annotation identified eight distinct subpopulations such as Astrocyte, Excitatory neuron (Excit neuron), Inhibitory neuron (Inhib neuron), Oligodendrocyte, Oligodendrocyte Precursor Cell (OPC), microglia, Endothelial cell, and Pericytes. (Fig. 1b). Notably, we observed a specific transcriptomic region where cells from HC and PD patients were highly interspersed, suggesting an intermediate or transitional phase rather than a definitive disease boundary. We classified this intermediate subpopulation as the BIF state.

**Fig. 1.**
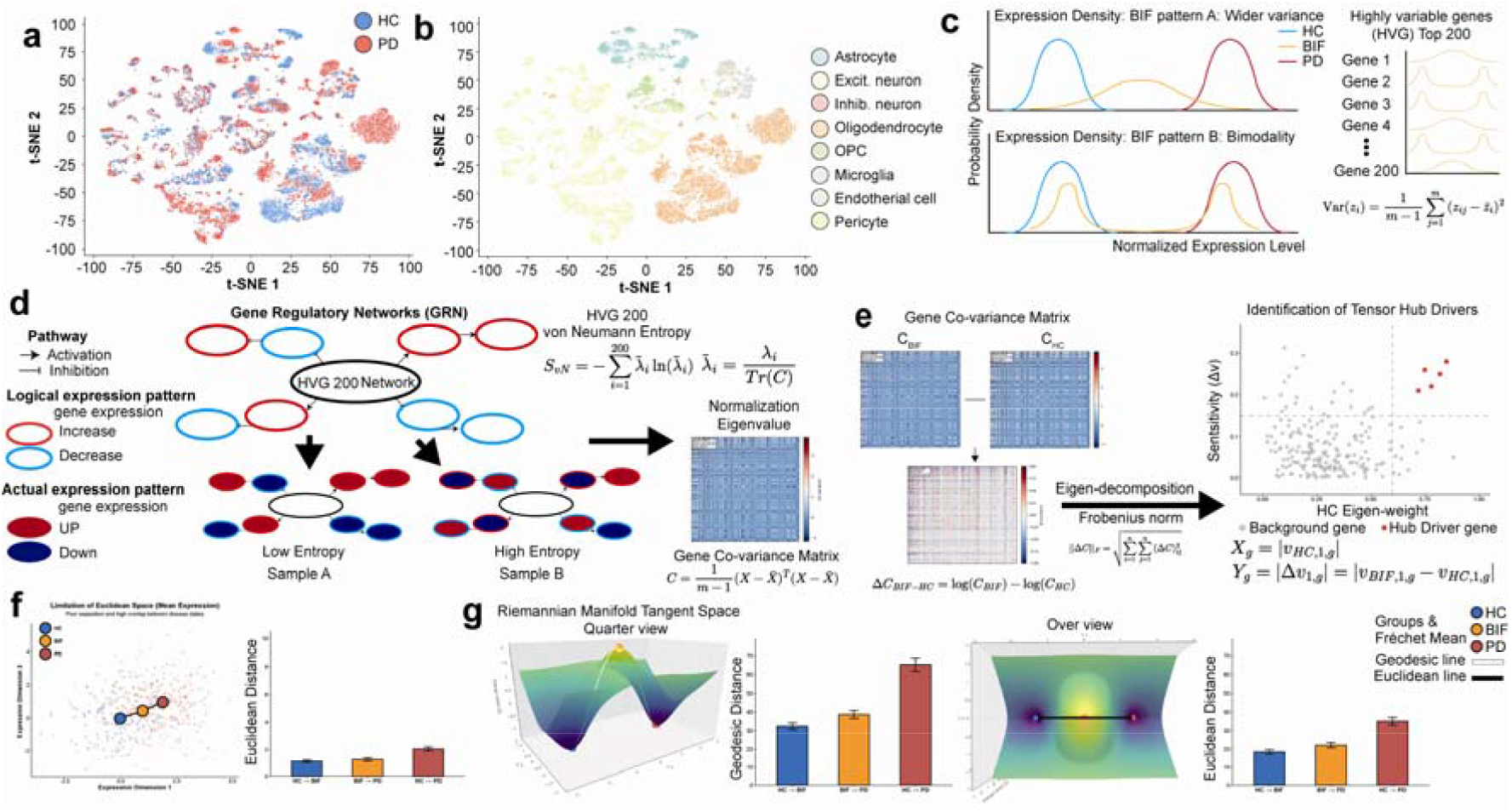
Identification of a disease-driving bifurcation state via Riemannian tensor analysis of single-cell transcriptomes. a, t-distributed stochastic neighbour embedding (t-SNE) visualization of the prefrontal cortex single-cell RNA-sequencing (scRNA-seq) dataset, capturing the broad cellular landscape across healthy control (HC) and Parkinson’s disease (PD) conditions. b, Cell-type annotations revealing eight distinct subpopulations, establishing the biological foundation for subsequent transcriptomic profiling. c, Mean-variance relationship demonstrating significant transcriptomic instability in a specific sample subset. These samples exhibited abnormally high standardized variance and bimodal expression density, warranting their classification as a transitional bifurcation (BIF) state. The top 200 highly variable genes (HVGs) were isolated for structural analysis. d, Methodological transition from mean-based expression to structural co-variance. The selected 200 HVGs were mapped into condition-specific gene co-variance matrices (*C*), representing complex transcriptomic rewiring. The von Neumann entropy analysis confirms maximal network disorder during the BIF state. e, Mathematical identification of hub driver genes. By computing the Frobenius norm on the differential matrix (Δ*C*_(BIF-HC)_), we extracted the extreme topological disturbers that drive the structural collapse from the healthy baseline. f, Limitations of traditional Euclidean space. Scatter plots and centroid distance quantification based on mean expression fail to definitively separate HC, BIF, and PD, yielding a structurally ambiguous and compressed trajectory. g, The Riemannian manifold workflow. Symmetric positive definite (SPD) covariance tensors were projected onto a non-Euclidean energy landscape. The Fréchet means for HC, BIF, and PD establish BIF as the high-energy saddle point. Geodesic distance measurements along the curved manifold conclusively distinguish the BIF state, proving its role as the critical tipping point in PD progression.

To investigate the molecular features distinguishing this BIF state, we examined the expression profiles of genes within this intermixed cluster compared to the healthy baseline. We found that the BIF state was characterized by pronounced transcriptomic instability, specifically exhibiting an abnormally widened variance or bimodal distribution in gene expression (Fig. 1c). Recognizing that this instability reflects systemic disorder, we isolated the top 200 highly variable genes (HVGs) showing these specific variance patterns for deeper structural analysis. This establishes that the initial departure from healthy homeostasis is not driven by a coordinated, unidirectional shift in gene expression, but rather by a massive, microscopic destabilization of transcriptional variance.

Traditional transcriptomic analyses often rely on mean expression levels, which may overlook complex systemic interactions. To address this, we evaluated whether the 200 HVGs followed logical regulatory pathways (low entropy) or exhibited random, disorganized expression (high entropy). This von Neumann entropy analysis confirmed that the network experiences maximal structural disorder during the BIF phase. To capture this systemic rewiring, we constructed condition-specific gene co-variance matrices (*C*) for the HC, BIF, and PD states (Fig. 1d).

We hypothesized that the critical transition is driven by specific genes disrupting the core network. By calculating the differential matrix (ΔC_BIF-HC_) and applying eigen-decomposition (measured via the Frobenius norm), we successfully identified the extreme topological disturbers defined here as hub driver genes that actively instigate the structural collapse from the healthy baseline (Fig. 1e).

The necessity for a non-linear geometric approach became evident when applying conventional analytical methods. Standard principal component analysis (PCA) based on differentially expressed genes (DEGs) failed to definitively separate the BIF group from HC and PD, yielding a compressed and structurally ambiguous trajectory (Fig. 1f). This limitation indicates that Euclidean geometry is insufficient to resolve the complex transition dynamics of neurodegeneration.

To overcome this, we projected the symmetric positive definite (SPD) covariance tensors onto a non-Euclidean Riemannian manifold (Fig. 1g). This geometric mapping revealed that the BIF group is positioned equidistantly between the HC and PD groups in the tensor space, functioning mathematically as a high-energy saddle point. Furthermore, by projecting this curved Riemannian manifold onto a 2D plane using a logarithmic mapping, we demonstrated that the Fréchet mean of the BIF state is distinctly separated from those of the HC and PD states. This geodesic distance measurement along the manifold conclusively identifies the BIF state not merely as an intermediate mix of cells, but as a mathematically defined critical tipping point in PD progression.

### Microscopic Variance Instability Reveals the Limitations of Conventional Transcriptomic Metrics

To further delineate the transitional dynamics between HC and PD, we mapped the integrated scRNA-seq data onto a comparative t-SNE projection (Fig. 2a). While distinct condition-specific clusters were evident, we observed a significant “mixed area” where HC and PD transcriptomes were highly overlapping. This continuous distribution highlights that the disease transition is not an abrupt shift, confounding clear macroscopic separation. When mapping this mixed area onto our previously defined cell-type annotations, we found that the overlap predominantly localized within excitatory neurons, oligodendrocytes, and astrocytes (Fig. 2b). This observation suggests that these specific glial and neuronal populations serve as the primary substrates for early structural rewiring prior to overt neurodegeneration.

**Fig. 2.**
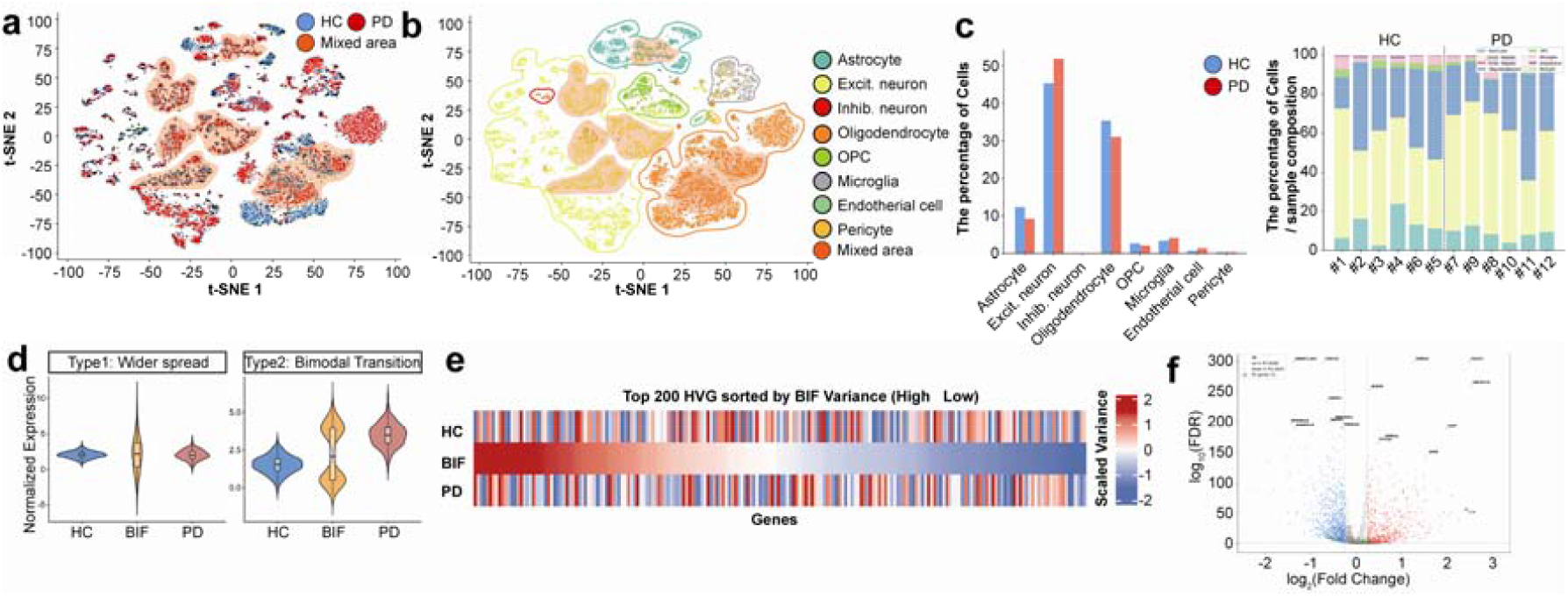
Microscopic variance instability underlying the bifurcation state reveals the limitations of conventional transcriptomic metrics. a, Comparative t-SNE projection of HC and PD single-cell transcriptomes. While distinct state-specific clusters exist, a significant “mixed area” highlights the continuous and overlapping nature of the disease transition, confounding macroscopic separation. b, Mapping of the mixed area onto cell-type annotations. The overlap predominantly localizes to excitatory neurons, oligodendrocytes, and astrocytes, suggesting these cell types are the primary substrates of early structural rewiring. c, Macroscopic cellular composition across states. Bar plots (left) indicate broad trends, such as the expansion of excitatory neurons, while individual sample compositions (right) demonstrate high inter-sample heterogeneity, rendering cellular proportions alone insufficient for defining the critical transition state. d, Representative expression distribution patterns underlying ‘critical slowing down.’ Genes at the bifurcation point exhibit extreme variance expansions rather than mean shifts, categorized into Type 1 (wider spread) and Type 2 (bimodal transition) patterns, indicating a loss of transcriptomic homeostasis. e, Variance heatmap identifying structural instigators. The heatmap displays the standardized variance (rather than mean expression) of the top 200 highly variable genes (HVGs), sorted in descending order based on their variance in the BIF state. f, A conventional differential expression (DEG) volcano plot comparing HC and PD fails to capture the core topological disturbers. By explicitly highlighting the top 200 HVGs onto this traditional volcano plot.

We next assessed whether broad cellular composition changes could define this transition. Quantitative comparison revealed significant shifts, such as an expansion of excitatory neurons (∼45% in HC to ∼52% in PD) and a contraction of astrocytes (∼12% to ∼9%) and oligodendrocytes (∼35% to ∼31%) (Fig. 2c, left). However, analysis of individual sample compositions demonstrated striking inter-sample heterogeneity. Notably, certain HC samples exhibited cellular proportions that closely resembled those of the PD state (Fig. 2c, right). This variability indicates that cellular proportions alone are insufficient for capturing the critical transition, prompting a deeper investigation into microscopic transcriptomic instability.

We hypothesized that the true hallmark of this intermediate state lies in the breakdown of transcriptomic homeostasis, manifesting as altered variance rather than shifts in mean expression. Indeed, transcriptional profiling of the samples exhibiting transitional characteristics revealed extreme variance expansions. These ‘critical slowing down’ patterns were broadly categorized into Type 1 (abnormally wide spread) and Type 2 (bimodal distribution) expression patterns (Fig. 2d). Crucially, because conventional differential gene expression (DEG) analyses rely on mean differences and low intra-group variance, these highly unstable genes yield p-values approaching 1, rendering them mathematically invisible to standard methodologies.

To systematically capture this instability, we calculated the standardized variance of all genes specifically within these transitional samples. We then isolated the top 200 HVGs, sorted in descending order of their variance, to serve as the foundation for our structural network analysis (Fig. 2e).

To explicitly demonstrate the limitations of traditional analytical frameworks, we projected these 200 HVGs onto a conventional DEG volcano plot comparing the broader HC and PD groups. As anticipated, traditional DEG analysis successfully identified canonical PD markers driven by mean expression shifts, such as the upregulation of *IL1B* (neuroinflammation) and heat shock proteins (*HSPA1A, HSPA1B*), alongside the downregulation of neuroprotective factors like *GDNF* and *SORCS2*. However, the 200 variance-driven HVGs identified in our BIF state consistently failed to meet conventional significance cut-offs (FDR < 0.05 and an absolute log2 fold change (|log2FC|) > 0.25) in the volcano plot (Fig. 2f). This critical finding proves that the core topological disturbers driving early disease pathogenesis are fundamentally masked when relying solely on mean-based differential expression, necessitating the application of higher-order tensor geometry.

### System-Wide Network Collapse and Geometric Identification of Topological Hub Drivers

Having established the limitations of mean-based differential expression, we applied Riemannian tensor geometry to capture the systemic rewiring of the 200 HVGs. We constructed condition-specific gene covariance matrices (C_HC_, C_BIF_, C_PD_) to represent the global network architecture for each clinical state (Fig. 3a). To conceptually understand the 200-dimensional eigendecomposition driving this network, we developed a 2D pedagogical geometric model (Fig. 3b). In this model, the principal eigenvector (v1) is constrained within a unit circle (|v|=1). The projection of v1 onto any specific gene expression axis yields its absolute eigen-weight (|v_1,i_|), mathematically quantifying that gene’s structural dominance within the macroscopic network. Applying this framework, we quantified the overall integrity and structural disorder of the network across disease progression (Fig. 3c). We discovered that the transition to the BIF state is characterized by a catastrophic, simultaneous collapse of total network energy (Trace) and principal module dominance (Max Eigenvalue, λ1). Concurrently, the von Neumann entropy (S) reached its maximum during the BIF state, before decreasing again in the PD state. This U-shaped entropic trajectory mathematically proves that the BIF phase is not merely an intermediate degradation, but a highly unstable ‘anarchic’ saddle point where healthy hubs collapse prior to the establishment of a new, low-entropy pathological order in PD. To identify the specific molecular drivers of this topological collapse, we traced the eigenvector trajectories across states (Fig. 3d, left). The line plot reveals a precipitous loss of eigen-weights for dominant hub genes during the HC-to-BIF transition. By plotting the initial structural importance (HC Eigen-weight) against this transition-induced instability (Rotational Sensitivity, Δv), we successfully identified the primary tensor hub drivers that uniquely trigger the breakdown of the healthy state (Fig. 3d, right). Specifically, this geometric analysis pinpointed top structural instigators *CNTNAP2, LHFPL3, TNR, GRIK1*, and *SORBS1* revealing that the macroscopic topological collapse is directly driven by the concurrent dismantling of core synaptic adhesion and electrophysiological scaffolds.

**Fig. 3.**
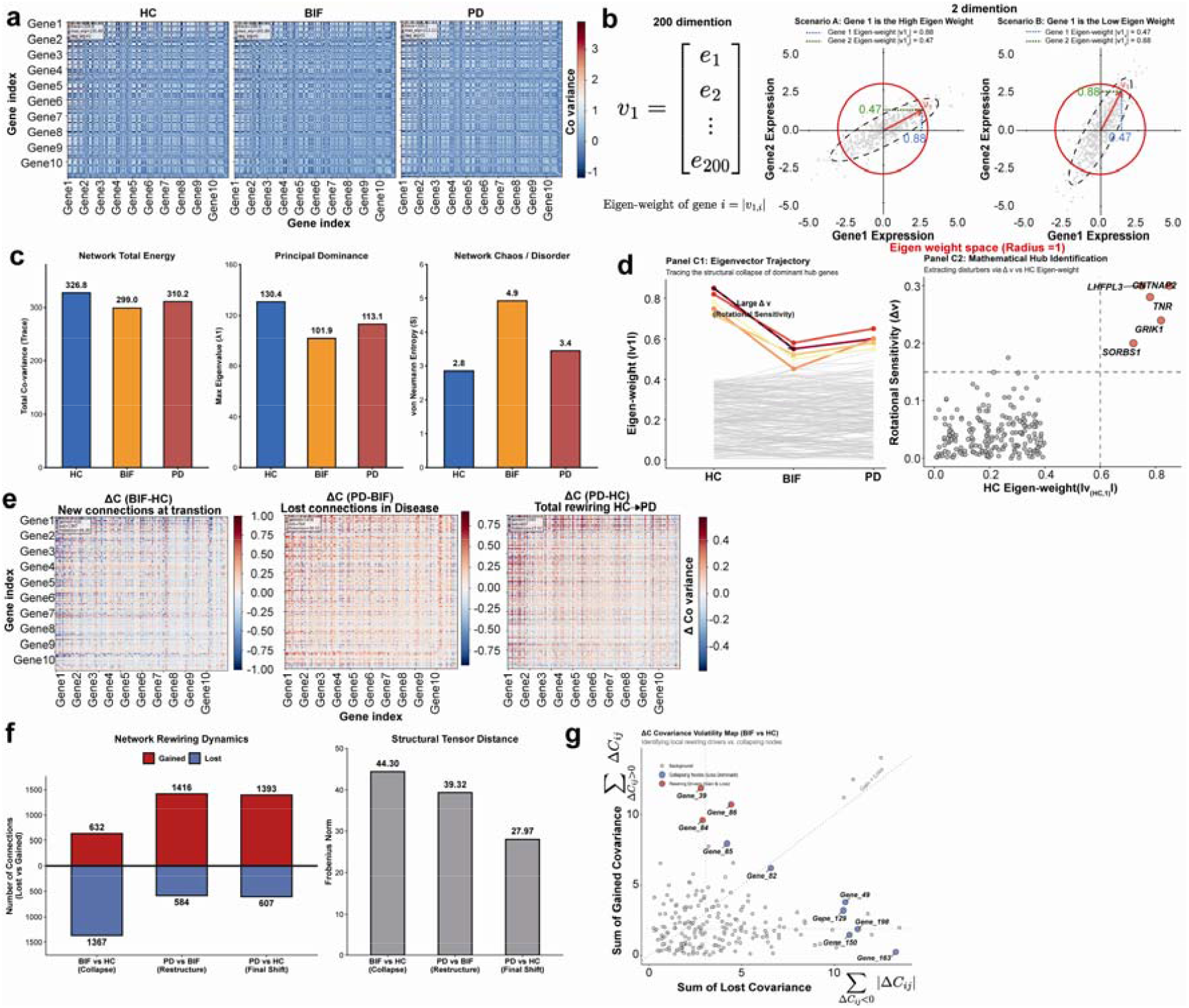
System-wide network collapse and geometric identification of topological hub drivers at the bifurcation state. a, Condition-specific gene co-variance matrices (C_HC_, C_BIF_, C_PD_) constructed from the top 200 highly variable genes (HVGs). b, Geometric projection of eigen-weight dynamics. To conceptually illustrate the 200-dimensional eigendecomposition, a 2D pedagogical model displays the geometric relationship between gene expression covariance, the principal eigenvector (red arrow, v_1_) constrained within a unit circle (Eigen-weight space, |v|=1). c, Quantification of network integrity and structural disorder. The critical transition is marked by a simultaneous collapse of total network energy (Trace) and principal module dominance (Max Eigenvalue, λ_1_), culminating in maximal von Neumann Entropy (S) during the BIF state. The projection of v_1_ onto the specific gene expression axis yields its absolute Eigen-weight (|v_1,i_|). d, Eigenvector trajectory tracing the structural collapse. The line plot demonstrates the precipitous loss of eigen-weights for dominant hub genes as the system transitions from HC to BIF, mathematically defining the rotational sensitivity (Δv) of each gene during the critical state (left), Mathematical identification of tensor hub drivers. A scatter plot mapping the initial structural importance (HC Eigen-weight, X-axis) against the transition-induced instability (Rotational Sensitivity, Δv, Y-axis) (right). e, Differential covariance matrices (ΔC_BIF-HC_, ΔC_PD-BIF_, ΔC_PD-HC_) visualizing the absolute changes in pairwise gene interactions across state transitions. f, Bar plots quantifying the global network rewiring. the massive proportion of lost connections versus gained connections during the initial HC-to-BIF transition, compared to the compensatory rewiring in the BIF-to-PD transition (left). the tensor distance (Frobenius norm, |ΔC|F), confirming the magnitude of structural disruption at each step (right). g, Covariance volatility map. A scatter plot characterizing the dual nature of network rewiring at the gene level by mapping the sum of lost covariance (X-axis) against the sum of gained covariance (Y-axis) for each gene during the BIF transition.

To further dissect the physical nature of this network rewiring, we computed the differential covariance matrices (ΔC_BIF-HC_, ΔC_PD-BIF_, ΔC_PD-HC_), which visualize the absolute changes in pairwise gene interactions (Fig. 3e). Quantification of these changes revealed a massive asymmetry in network dynamics (Fig. 3f, left). To rigorously filter background noise and isolate biologically meaningful rewiring events, we applied a predefined significance threshold - retaining only the top 10% of varying edges / edges exceeding a Z-score of 2.0] to the differential matrices. Under this criterion, the initial transition into the BIF state (ΔC_BIF-HC_) was overwhelmingly defined by structural loss, with 1,367 lost connections compared to only 632 gained connections. This net deficit of 735 connections provides direct topological proof that the BIF saddle point represents an extreme dismantling of the healthy transcriptomic architecture, rather than a mere shift in cellular state. Conversely, the subsequent progression from BIF to PD was dominated by connection gains, reflecting the establishment of a new pathological network. The magnitude of this structural disruption was confirmed by the tensor distance (Frobenius norm, ||ΔC||_F_), which peaked dramatically at the BIF-HC transition (44.30), significantly exceeding both the PD-BIF (39.32) and overall PD-HC (27.97) distances (Fig. 3f, right).

Finally, to characterize this dual nature of network rewiring at the single-gene level, we constructed a Covariance Volatility Map (Fig. 3g). By mapping the sum of lost covariance against the sum of gained covariance for each gene during the BIF transition, we identified specific hub drivers that actively dismantle healthy architecture while simultaneously orchestrating the formation of pathological connections.

### Riemannian Tensor Geometry Identifies the Bifurcation State as a Critical Topological Saddle Point

To precisely define the topological nature of the BIF state, we compared conventional analytical methods with Riemannian tensor geometry. A conventional PCA, which relies on mean expression levels within a flat Euclidean space, failed to structurally differentiate the disease stages, resulting in overlapping regions where the HC, BIF, and PD states could not be clearly resolved (**Fig. 4a**). This highlights the fundamental limitation of Euclidean space in capturing complex, non-linear disease transitions.

**Fig. 4.**
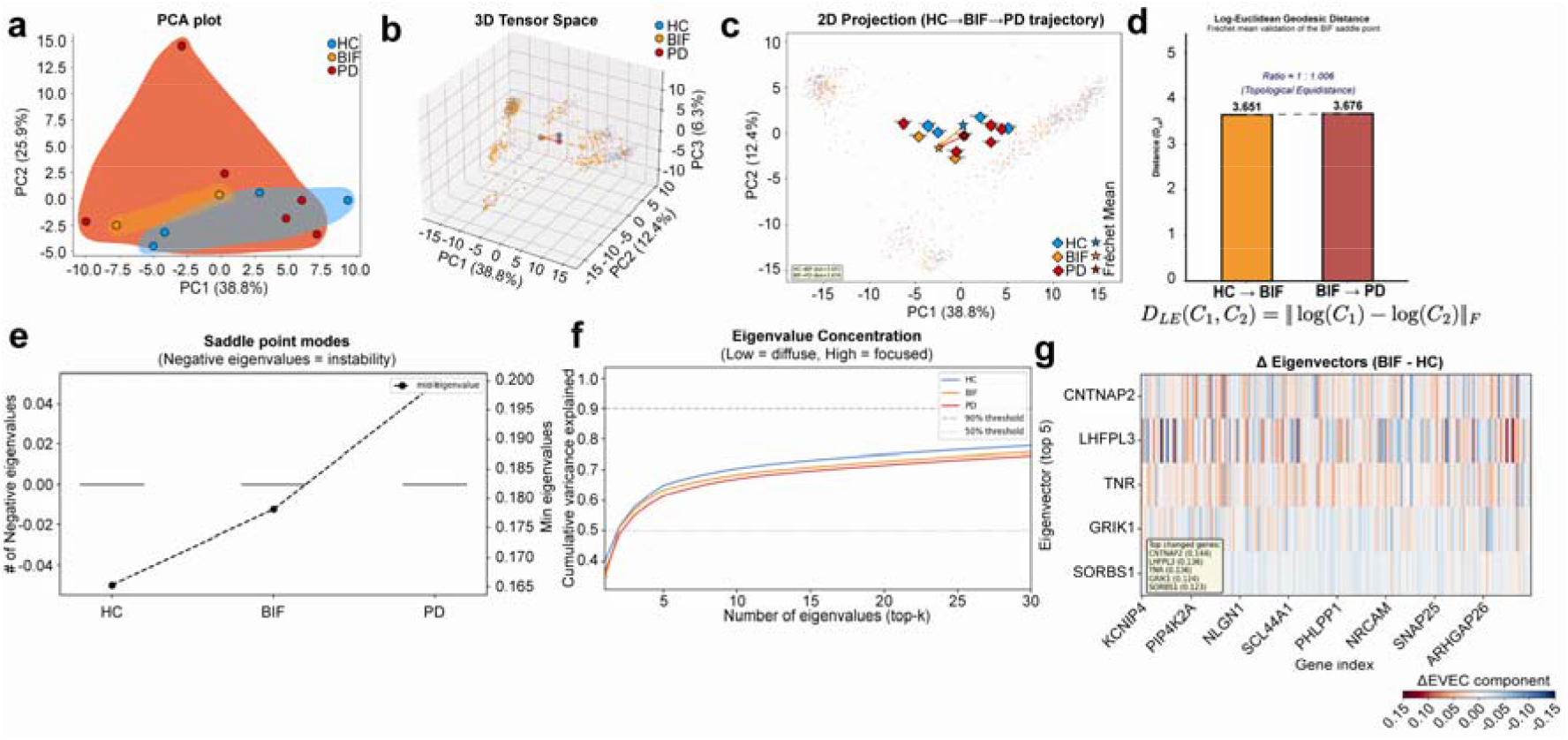
Riemannian tensor geometry identifies the bifurcation state as a critical topological saddle point of synaptic network collapse. a, Conventional principal component analysis (PCA) plot of gene expression from differentially expressed genes (DEGs). b, Three-dimensional Riemannian tensor space representation constructed from the covariance matrices of the top 200 highly variable genes (HVGs). **c**, Two-dimensional Log-Euclidean projection (Tensor PCA) of the Riemannian space. **d**, Bar plot depicting the log-Euclidean geodesic distance (D_LE_) between the Fréchet means of each state. **e**, Line graph showing the progressive increase of the minimum eigenvalue (λ_min_). **f**, Cumulative variance explained by the top-*k* eigenvalues. **g**, Heatmap displaying the delta eigenvector (ΔV) components of the top 5 rotational sensitivity hubs (*CNTNAP2, LHFPL3, TNR, GRIK1, SORBS1*) alongside 8 key functionally enriched synaptic genes.

To overcome this, we constructed a three-dimensional Riemannian tensor space using the covariance matrices of the top 200 HVGs (**Fig. 4b**). In this non-Euclidean energy landscape, the samples from each group along with their respective Fréchet means (geometric centroids) were distinctly segregated, revealing a highly structured trajectory. Because interpreting a 3D tensor manifold can be mathematically abstract, we projected this manifold onto a two-dimensional Log-Euclidean space (Tensor PCA) while preserving the intrinsic geometric relationships (**Fig. 4c**). This projection clearly plotted all 12 individual samples and their corresponding Fréchet means, illustrating a defined transition path from HC to BIF (orange arrow) and subsequently from BIF to PD (red arrow). Crucially, quantifying the Log-Euclidean geodesic distance (*D*_LE_) between these Fréchet means provided definitive mathematical evidence for the critical transition. The distance from the HC centroid to the BIF centroid was 3.651, nearly identical to the distance from the BIF centroid to the PD centroid (3.676) (**Fig. 4d**). This remarkable geometric equidistance (distance ratio = 0.993) mathematically proves that the BIF state operates as a central saddle point, situated exactly at the energetic threshold between the healthy and pathological attractors.

We further confirmed this topological shift through eigenvalue spectrum analysis of the gene co-variance tensors. While the maximum eigenvalue achieved a minimum at the BIF state (101.9, reflecting the transient collapse of global dominance), the minimum eigenvalue (λ_min_) exhibited a continuous, monotonic increase from HC (0.165) to BIF (0.180) to PD (0.195) (**Fig. 4e**). This steady elevation of λ_min_ signifies a progressive stiffening of the energy landscape and the irreversible loss of functional network isolation, as isolated nodes increasingly form ubiquitous, noisy connections due to chronic cellular stress.

This disruption of centralized control was corroborated by the cumulative variance explained by the top-*k* eigenvalues (**Fig. 4f**). The highly focused curve of the HC state, where a few dominant modules efficiently govern the global system, shifted significantly downward in the BIF state, reflecting a ‘low diffuse’ pattern. This downward shift mathematically demonstrates the fragmentation of the centralized network leadership, pushing the system’s variance toward the highly shattered trajectory observed in PD.

Finally, to bridge these macroscopic geometric findings with specific molecular pathology, we evaluated the rotational sensitivity (ΔV) driving this tensor space rotation during the HC → BIF transition (**Fig. 4g**). A heatmap of the ΔV components identified five primary tensor hub drivers (*CNTNAP2, LHFPL3, TNR, GRIK1, SORBS1*) alongside eight key functionally enriched synaptic genes. The abrupt loss of eigen-weights in these specific structural and presynaptic hubs at the BIF state implicates the dismantling of synaptic architecture as the earliest molecular event precipitating the macroscopic transition into PD.

## Discussion

In this study, we established a novel mathematical framework utilizing Riemannian tensor geometry to decode the transcriptomic collapse underlying the progression of PD. Our analytical pipeline successfully identified a critical transitional phase termed the BIF state that operates as an energetic saddle point between the HC and PD attractors. By moving beyond conventional expression-level metrics and examining macroscopic covariance trajectories, our approach mathematically isolated the BIF state with high topological precision and identified a completely distinct set of early-stage structural instigators, including crucial synaptic hubs such as *CNTNAP2*^*27*^, *LHFPL3*^*28*^, *TNR*^*29*^, *GRIK1*^*30*^, and *SORBS1*^*31*^.

The necessity for such a paradigm shift in analytical methodology is deeply rooted in the current stagnation of therapeutic target discovery. Over the past decade, extensive meta-analyses integrating RNA-seq, proteomics, and metabolomics have sought to elucidate the pathological hallmarks of various neurodegenerative diseases ^32-34^. While these comprehensive molecular profiles have yielded numerous candidate markers based primarily on differential gene expression ^32^, they have largely fallen short of translating into effective disease-modifying therapies in clinical settings. This persistent translational gap suggests a fundamental limitation in how we traditionally define and target the temporal phases of neurodegeneration, particularly regarding the events that occur strictly prior to irreversible cellular damage.

The pathogenesis of neurodegenerative diseases is driven by the progressive accumulation of toxic aggregates such as alpha-synuclein, amyloid-beta, hyperphosphorylated tau, and mutant huntingtin which profoundly disrupt the homeostatic balance of both neuronal and glial populations ^35, 36^. Crucially, these cell types exhibit highly dynamic state transitions in response to toxic microenvironments ^37^. Astrocytes, for instance, can shift seamlessly between protective, harmful, and chronic reactive states ^38^, while microglia exhibit a complex spectrum of activation profiles whose chronic stages differ fundamentally from their initial responses ^39^. Similarly, neurons undergo extensive synaptic deterioration and structural rewiring long before the onset of irreversible apoptosis ^40, 41^. This highly dynamic nature of cellular responses underscores the critical need to intercept the disease trajectory during its earliest, reversible phases, rather than attempting to rescue terminally exhausted neural networks.

Recognizing the paramount importance of early intervention, numerous studies have attempted to delineate the pre-disease boundary to establish effective therapeutic strategies ^1, 42^. In clinical settings, stage-specific scoring systems are widely utilized to track pathological progression by integrating cognitive evaluations, motor behavioral assessments, and histological criteria. For instance, the MDS-UPDRS (Movement Disorder Society-Sponsored Revision of the Unified Parkinson’s Disease Rating Scale) and the Hoehn and Yahr scale provide standardized clinical evaluations of motor and non-motor deficits in PD ^43, 44^. Complementing these behavioral metrics, Braak staging offers a definitive post-mortem pathological classification based on the ascending anatomical spread of alpha-synuclein Lewy pathology ^45^. However, because these established metrics inherently rely on the manifestation of overt neurological symptoms or macroscopic protein aggregation, they are fundamentally limited in their ability to pinpoint the pre-symptomatic, molecular tipping points that precede irreversible network fragmentation. Concurrently, basic research frequently employs presymptomatic animal models for example, analyzing transcriptomic changes in younger mice prior to the onset of overt behavioral deficits or severe histological pathology to identify putative early targets ^46, 47^. Despite these extensive efforts, defining a precise biological boundary that separates healthy aging from the irreversible commitment to disease remains highly elusive. Consequently, the field still lacks a definitive methodology to isolate the ‘tipping point’ before overt clinical symptoms manifest.

Perhaps the most profound limitation of conventional transcriptomic analysis lies in its inherent statistical architecture, which fundamentally prioritizes mean-based differences while heavily penalizing variance ^**48**, **49**^. Standard DEG frameworks ranging from foundational statistical tests (e.g., Student’s *t*-test, Wilcoxon rank-sum test)^50^ to widely adopted bioinformatics pipelines (e.g., DESeq2, edgeR) evaluate mean expression shifts relative to intra-group dispersion ^51, 52^. Whether structured as a direct denominator in a classical test statistic or as an overdispersion penalty within a negative binomial model, the mathematical consequence remains identical. Consequently, when a gene exhibits massive variance expansion or a bimodal distribution which are, in fact, crucial biological hallmarks of a system approaching a critical tipping point this dispersion inflates disproportionately ^53, 54^. This structural artifact mathematically forces the test significance to converge toward zero, invariably yielding *p*-values approaching 1 ^48^. Through this mathematical penalty, conventional algorithms inherently discard the most dynamic, state-transitioning genes as non-significant biological noise. Consequently, these genes are erroneously discarded as non-significant biological noise. However, during our analysis of clinical scRNA-seq datasets, we observed that certain samples despite being clinically classified as healthy controls displayed precisely this erratic transcriptomic behavior. Recognizing that such variance expansion is a classic thermodynamic hallmark of a complex system approaching a critical tipping point, we redefined these fluctuations not as noise, but as the earliest indicators of lost transcriptomic homeostasis. By isolating the top 200 HVGs and characterizing the BIF state based on this variance instability, our approach successfully illuminates the critical temporal blind spot entirely missed by traditional DEG profiling.

Crucially, gene expression does not alter in isolation; it operates within highly coordinated gene regulatory networks (GRNs) governed by strict logical pathways ^55, 56^. A healthy system maintained by these robust GRNs exhibits low structural entropy, whereas the stochastic, disorganized transcription characteristic of the BIF state drives network entropy to its mathematical maximum ^25^. Through eigen-decomposition of these 200-dimensional matrices, we computed the total network energy (Trace) and the specific eigen-weight of each gene ^57^, quantitatively defining the structural dominance a single node exerts over the remaining 199 interactions ^58, 59^. While tensor calculus and Riemannian geometry have profound historical roots in theoretical physics and macroscopic neuroimaging (e.g., Diffusion Tensor Imaging) ^60^, applying these continuous topological mappings to the sheer dimensionality of single-cell transcriptomic networks was previously computationally prohibitive ^4, 61^. It is the recent advent of high-performance, GPU-accelerated tensor computing ecosystems initially spearheaded by the machine learning community that has finally made the dynamic geometric decoding of such massive biological systems feasible, positioning this methodology at the vanguard of modern computational biology ^62, 63^.

Furthermore, conventional analytical strategies that stratify disease progression into discrete, static clusters yield only an isolated snapshot, effectively masking the fluid continuum of biological transitions ^7^. From the perspective of network medicine, neurodegeneration arises not from an isolated molecular defect at a single time point, but from the dynamic, longitudinal perturbation of complex regulatory networks; therefore, relying on such static cross-sectional analyses is fundamentally insufficient for uncovering true therapeutic targets ^64^. The paramount value of our Riemannian tensor framework lies in its capacity to dynamically track macroscopic network volatility across continuous state transitions (HC → BIF, BIF → PD, and HC → PD). By quantifying the differential covariance (Δ*C*) and the degree of eigenspace rotation (Δ*v*) for each gene across these distinct phases, we can trace exactly how the network is dismantled and rewired. Consequently, genes demonstrating the most dramatic eigen-weight fluctuations during the initial HC → BIF transition are not merely downstream markers of disease; they are the fundamental ‘key switches’ that actively orchestrate the entry into the neurodegenerative trajectory, serving as the most optimal targets for early intervention.

The most compelling mathematical evidence supporting the critical nature of the BIF state emerged from our topological analysis within the Riemannian manifold ^20^. While conventional Euclidean PCA struggled to isolate the disease groups ^22^, mapping the covariance tensors onto a three-dimensional Riemannian space definitively segregated the HC, BIF, and PD samples. Strikingly, when measuring the log-Euclidean geodesic distances between the Fréchet means of these respective clusters, we discovered a near-perfect equidistance (ratio almost 1) between the HC-to-BIF and BIF-to-PD trajectories. This geometric symmetry mathematically dictates that the BIF state is not merely an intermediate degradation phase, but a structural ‘saddle point’ operating at the highest energy level between two distinct attractors ^65-67^. From a biological and clinical perspective, this topological positioning is of paramount importance. A saddle point represents a highly unstable, yet transiently balanced state where the network has lost its healthy architecture but has not yet fully committed to the pathological attractor ^65, 66^. Once the system undergoes a complete phase shift into the PD state, the newly established pathogenic network characterized by extensive synaptic loss and chronic glial reactivity becomes structurally entrenched and largely irreversible. Therefore, the BIF state signifies the ultimate window of therapeutic reversibility. By leveraging our tensor-derived hub drivers (*CNTNAP2, GRIK1*, etc.) to selectively target the network while it resides at this tipping point, we propose a novel strategic paradigm designed to arrest neurodegeneration before the irreversible commitment to disease pathology occurs.

While our Riemannian tensor framework provides profound topological insights into the transcriptomic collapse at the BIF state, we acknowledge the limitations regarding the data volume utilized in this study, which was based on a cohort of 12 clinical scRNA-seq samples. The predictive power and robustness of mathematical modeling inherently benefit from large-scale validation. Therefore, our future studies will aim to leverage massive clinical and multi-omics datasets, such as those provided by the ROSMAP and the Dementias Platform UK (DPUK). By applying our variance-based Riemannian tensor methodology to these extensive, well-characterized clinical cohorts, we seek to validate the universal existence of the BIF saddle point across diverse patient populations. Ultimately, this expanded approach will enhance the translational value of our analytical framework, paving the way for the development of highly customized, BIF-stage-specific therapeutic strategies that can halt neurodegeneration before irreversible network fragmentation occurs.

## Conclusion

In summary, by employing a Riemannian tensor framework that transcends the limitations of conventional mean-based transcriptomics, we have successfully identified the ‘BIF state’ a topological saddle point along the trajectory of PD progression. Driven by catastrophic network rewiring and the collapse of core synaptic hubs, this BIF state is not merely an intermediate phase of degeneration; rather, it represents the ‘ultimate window of biological reversibility’ before the macroscopic system irreversibly fragments into a pathological attractor. Ultimately, the geometric blueprint provided by this study shifts the therapeutic paradigm from attempting to repair already decimated networks toward stabilizing this critical tipping point. This conceptual pivot paves the way for preemptive, disease-modifying strategies capable of intercepting the neurodegenerative trajectory at its earliest stage, long before the pathology becomes entrenched.

## Methods

### Data Acquisition and Variance-Preserving Preprocessing

Raw scRNA-seq data, spanning 12 samples from the human prefrontal cortex of PD patients and HC (GEO accession: GSE202210) ^26^, were processed using the Cell Ranger pipeline (v6.1.1, 10x Genomics) to generate initial gene expression matrices. Downstream quality control (QC) and preprocessing were conducted using Seurat (v4.3.0) in R (v4.2.1) ^68, 69^. To establish a rigorous baseline of cellular viability, we applied stringent QC thresholds: cells expressing fewer than 200 or more than 6,000 genes, as well as those with a mitochondrial read fraction exceeding 20%, were strictly excluded.

Following basic QC, it was imperative to decouple true biological volatility from technical sequencing depth without mathematically penalizing the macroscopic variance essential for tensor construction. Standard log-normalization pipelines inherently distort high-variance signals. To circumvent this, we employed the SCTransform framework to compute analytic Pearson residuals ^70^. Because these residuals explicitly quantify the deviation of each gene from its expected baseline model, they effectively regress out technical noise while perfectly preserving the extreme heteroscedasticity and bimodal distributional structures of HVGs. By deliberately utilizing these variance-preserved residuals, we ensured that the “thermodynamic engine” of the pre-disease state was mathematically conserved for the subsequent formulation of SPD covariance tensors.

To eliminate technical batch effects across the 12 independent samples while rigorously preserving intrinsic biological coordination, we employed the Harmony algorithm (v0.1.1) ^71^. Harmony aligns cellular manifolds by iteratively adjusting expression profiles without over-correcting local variance ^72^. The integrated dataset was then subjected to PCA. The top 50 principal components (PCs) were utilized for initial dimensionality reduction and t-SNE visualization ^73^, confirming the successful global integration of the data prior to the localized extraction of covariance tensors.

### Isolation of high variant genes

Isolation of Thermodynamic Hub Candidates via Variance InstabilityFollowing the extraction of analytic Pearson residuals via the SCTransform framework, which effectively neutralizes technical mean-variance dependencies, we systematically quantified the intrinsic biological volatility of each gene. Let 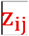 denote the normalized residual expression of gene 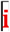 in cell 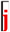. To capture the destabilization of the transcriptomic network, the standardized variance for each gene across the cellular population (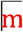 cells) was rigorously computed as:

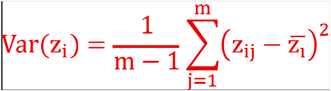

where 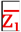 is the mean residual expression of gene 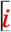. As a complex biological system approaches a critical transition (the BIF state), it exhibits a thermodynamic phenomenon known as ‘critical slowing down.’ In our single-cell framework, this instability manifests either as a massive uniform expansion of variance (wider spread) or the splitting of cellular states into bimodal expression landscapes. Because both patterns mathematically drive the 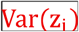 to extreme maximums, this formula serves as the exact quantitative metric for identifying network instability. By ranking the entire transcriptome based on this variance metric, we isolated the top 200 HVGs. These 200 genes, representing the most structurally volatile elements dismantling the pre-disease state, were subsequently defined as the fundamental coordinate axes 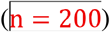 for the construction of our high-dimensional covariance tensors.

### Identification and annotation of cell types

To delineate the macroscopic phenotypic landscape of the prefrontal cortex ecosystem with strict objectivity, we eschewed traditional heuristic, marker-based clustering. Instead, we implemented a mathematically robust, reference-based mapping strategy. As our canonical manifold, we anchored our query data to the Human Multiple Cortical Areas SMART-seq transcriptomic taxonomy generated by the Allen Institute for Brain Science ^74^. Because this reference atlas is constructed from ultra-deep, full-length transcript sequencing, it provides a structurally flawless topological baseline. This approach enables the precise assignment of cell identities based on global transcriptomic similarity and high-resolution structural homology, rather than relying on the dimensional constraints of a limited subset of marker genes.

To project our query dataset onto this canonical reference space, we employed an anchor-based integration framework implemented in Seurat. Specifically, homologous coordinates—or mutual nearest neighbors—between the reference and query manifolds were identified (FindTransferAnchors) ^75^. This foundational alignment facilitated the robust propagation of cell-type identities from the HCA reference to our query dataset (TransferData) ^68^. By leveraging the shared transcriptional structure in a low-dimensional space, predicted identities were systematically consolidated into eight fundamental architectural lineages: Excitatory Neurons, Inhibitory Neurons, Astrocytes, Oligodendrocytes, OPCs, Microglia, Endothelial cells, and Pericytes.

### Geometric Construction of Covariance Tensors and Spectral Deconstruction

To characterize higher-order transcriptomic organization beyond mean expression levels, gene–gene covariance matrices were constructed for each sample. This approach captures coordinated variation between genes across single cells, allowing the representation of transcriptional states in terms of gene–gene relationships rather than individual gene abundances.

A subset of HVGs with consistent detection across samples was selected to ensure robust and stable estimation of covariance structure. For each sample, let 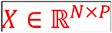 denote the gene expression matrix, where 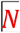 is the number of cells 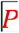 and is the number of selected genes. The covariance matrix 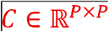 was defined as:

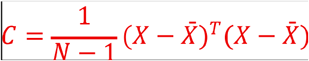

where 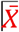 represents the mean expression vector across cells.

Each element 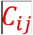 quantifies the degree to which genes 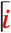 and 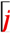 vary together across cells. Positive values indicate coordinated activation, whereas negative values reflect opposing patterns of expression. This formulation enables the characterization of transcriptional programs as structured interaction networks between genes.

### Covariance Rewiring and Spectral Deconstruction

To extract the fundamental structural components of the GRN, spectral eigendecomposition was applied to each tensor:

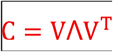

Here, 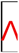 is a diagonal matrix of eigenvalues 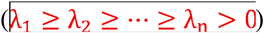, and 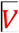 is an orthogonal matrix of eigenvectors 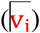. In our biological framework, each eigenvector represents an independent, orthogonal mode of gene co-expression, while its corresponding eigenvalue 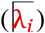 denotes the magnitude of variance captured by that module. The eigenvalue spectrum provides direct insight into how transcriptomic variance is distributed. For example, large dominant eigenvalues indicate highly coordinated variation along specific gene modules, whereas an increased dispersion of eigenvalues strictly reflects the structural instability and fragmentation of transcriptional programs characteristic of the bifurcation transition. This formulation enables the precise identification of dominant transcriptomic programs driving the covariance rewiring between HC and PD states.

### Covariance Rewiring Extraction of Tensor Hub Drivers via Eigen-weights

To pinpoint the exact genetic effectors driving the transition, we focused on the principal eigenvector 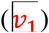 corresponding to the largest eigenvalue 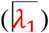, representing the primary axis of maximal transcriptomic variance. In our 200-dimensional covariance space, 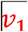 is defined as a column vector:

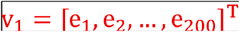

By mathematical definition, eigenvectors extracted from SPD matrices are strictly constrained to a unit hypersphere, satisfying the 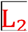 norm condition 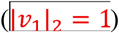. Because the total length of the vector is fixed, the absolute magnitude of each constituent coordinate precisely quantifies the proportional directional contribution of that specific gene to the dominant network motif. Accordingly, the ‘Eigen-weight’ of individual gene 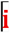 was formulated as the absolute value of its corresponding vector element:

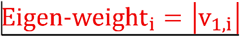

### Quantification of Network Disorder via von Neumann Entropy

As biological systems approach a critical transition (the bifurcation or BIF state), the coordination within the GRN typically undergoes severe destabilization. To quantitatively capture this thermodynamic hallmark, we computed the von Neumann entropy 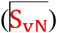 of the 200-HVG network.

Originating from quantum statistical mechanics, von Neumann entropy provides a rigorous measure of structural disorder within SPD matrices. We first normalized the covariance tensor to construct a transcriptomic density matrix 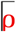:

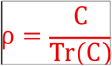

The network entropy was then calculated using the normalized eigenvalues 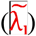:

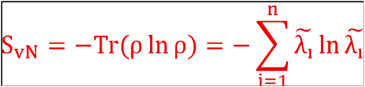

A localized peak in 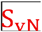 indicates an equipartition of variance across multiple eigenvectors—a flattened regulatory energy landscape characteristic of a highly volatile, transitional BIF state immediately preceding the commitment to the pathological PD attractor.

### Differential Tensor Calculus and Delta Covariance Networks

To decode the directional flow of transcriptomic rewiring between healthy and pathological states, we formulated a differential tensor calculus framework. Direct Euclidean subtraction of two covariance matrices 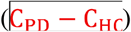 produces geometric artifacts. Therefore, the precise topological difference, or delta covariance 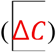, must be computed within the flat tangent space (Lie algebra) of the Riemannian manifold.

Using the HC state as the structural baseline 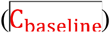, the transcriptomic rewiring toward a target state (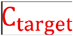, e.g., the BIF or PD state) was defined as:

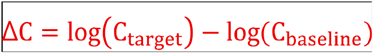

Mathematically, the resulting 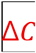 matrix is symmetric but no longer constrained to be positive definite. To comprehensively quantify the magnitude of this transition, the structural change was summarized using the Frobenius norm 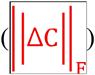. For our 200-dimensional gene network 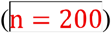, this norm was precisely computed as the square root of the absolute sum of all squared elements within the tangent space:

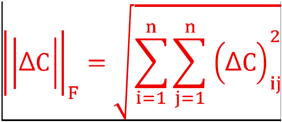

This measure elegantly aggregates all pairwise changes in gene-gene relationships into a single scalar quantity, reflecting the overall extent of structural reorganization. This relaxation inherently allows its eigenvalue spectrum to span both real positive and negative domains, providing a mathematically exact foundation for segregating emergent network formations from disintegrating homeostatic structures.

### Spectral Segregation of Gained and Lost Regulatory Pathways

To biologically interpret the 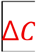 matrix, we performed spectral eigendecomposition:

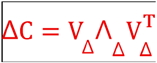

where 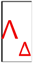 is the diagonal matrix of differential eigenvalues and 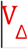 contains the corresponding differential eigenvectors. In this differential framework, the polarity of the eigenvalue explicitly dictates the biological fate of the regulatory module:

- **Gained Pathways (**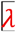 **> 0):** Eigenvectors associated with positive eigenvalues represent coordinated transcriptomic modules whose covariance variance significantly increased in the target state. These denote the ‘Gain-of-function’ rewiring—typically emergent pathological networks or compensatory stress responses.
- **Lost Pathways (**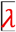 **< 0):** Eigenvectors associated with negative eigenvalues represent modules whose covariance was structurally dismantled. These denote the ‘Loss-of-function’ rewiring, signifying the collapse of healthy homeostatic interactions.

To extract the specific genetic drivers, the eigen-weights 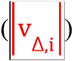 were quantified for the principal eigenvectors of both the extreme positive (maximum gain) and extreme negative (maximum loss) spectra. The top-ranking tensor hub drivers from these respective axes were systematically isolated. Finally, to translate these geometric hubs into functional biological contexts, standard over-representation analysis (ORA) and pathway enrichment were conducted on the extracted ‘Gained’ and ‘Lost’ gene sets using [사용한 패키지 예: clusterProfiler (v4.6.0)], referencing canonical databases including Gene Ontology (GO) and Kyoto Encyclopedia of Genes and Genomes (KEGG).

### Pseudobulk Baseline (Scalar PCA)

To capture baseline sample-level transcriptomic variation while mitigating cell-level stochastic sparsity, we first generated pseudobulk expression profiles by aggregating read counts across all cells within each defined state. By mapping these normalized scalar profiles via standard PCA, we established a Euclidean low-dimensional embedding. While this traditional pseudobulk PCA preserves global transcriptional trends and mean-level shifts between HC and PD states, it is mathematically blind to the higher-order structural rewiring of the gene regulatory network. Thus, it serves as a scalar baseline against which our covariance-based topological framework is evaluated.

### Riemannian Tangent Space PCA (tPCA)

To decode the hidden structural trajectory of the disease, dimensionality reduction must be performed on the network architectures themselves. Because the covariance tensors 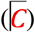 lie on the curved 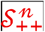 manifold, applying standard PCA directly produces severe geometric distortions. Therefore, we projected the tensors into a flat tangent space at the identity matrix using the Log-Euclidean mapping framework. Principal component analysis was subsequently performed on these vectorized tangent space embeddings (tPCA). This geometry-aware projection translates the high-dimensional covariance dynamics into a strictly accurate 2D topological coordinate system, enabling the direct visualization of structural relationships between transcriptomic states without Euclidean deformation.

### Fréchet Mean Calculation

Within this exact topological 2D space, the definitive biological attractors—the baseline HC and the irreversible pathological state (PD)—cannot be represented by simple arithmetic means. Instead, we computed their theoretical geographic centers as Fréchet means 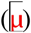. The Fréchet mean represents the geometric centroid on a Riemannian manifold, mathematically defined as the tensor that minimizes the sum of squared Log-Euclidean distances 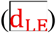 across all samples within a given biological group:

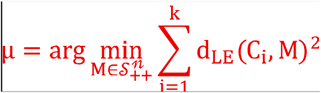

We computed the specific centroids for both the healthy attractor 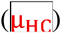 and the pathological attractor 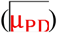, establishing the absolute anchor points of the disease trajectory.

### Topological and Thermodynamic Validation of the Saddle Point

TheBIF state was rigorously validated as the critical tipping point—the saddle point—of the disease progression. We operationally defined and mathematically verified this saddle point through a dual topological and thermodynamic validation:

1. Topological Equidistance: In the Riemannian embedding space, the BIF state must act as the exact geometric pivot. We computed the Log-Euclidean distances from the BIF tensor 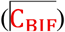 to both respective attractors, verifying that it occupies the structurally intermediate coordinate where the distance ratio approaches unity:

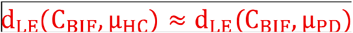
2. Thermodynamic Maximum: This exact topologically equidistant coordinate must strictly coincide with the peak of internal network volatility. We cross-validated that the 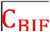 tensor exhibits the absolute maximum von Neumann entropy 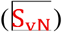 and the peak Frobenius magnitude of covariance rewiring 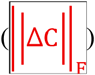.

By proving the convergence of these two independent mathematical criteria—exact geometric equidistance between Fréchet means and peak thermodynamic entropy—we robustly validated the BIF state not merely as an intermediate phase, but as the fundamental, highly volatile topological saddle point immediately preceding irreversible commitment to the Parkinsonian attractor.

### Dual-Metric Riemannian Framework and Topological Distances

To comprehensively characterize the dynamic trajectory of disease progression and pinpoint the critical tipping point of the network, we employed a complementary dual-metric Riemannian framework. Because covariance matrices 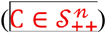 lie on a highly curved non-Euclidean manifold, direct Euclidean comparisons inherently fail to capture higher-order structural differences. To overcome this, we integrated two distinct but complementary non-Euclidean metrics.

### Affine-Invariant Riemannian Metric (AIRM) for Intrinsic Distortion

First, to strictly quantify the absolute magnitude of covariance rewiring between distinct transcriptomic states, we utilized the affine-invariant Riemannian metric (AIRM):

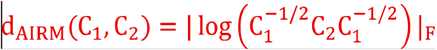

This formulation measures the true geodesic distance between covariance matrices directly on the curved manifold. It captures intrinsic structural differences while rigorously accounting for transcriptomic scaling and correlation patterns. In this framework, two transitional states are considered biologically similar only if their overall covariance structures can be transformed into each other with minimal geometric distortion.

### Log-Euclidean Metric (LEM) for Tangent Space Trajectory

While AIRM provides the gold standard for pairwise geodesic distance, performing complex statistical operations (e.g., spatial trajectory embedding, Fréchet mean estimation) directly on the curved manifold can introduce computational instability. Thus, for topological trajectory mapping, we complementarily mapped the covariance tensors to a flat tangent space (Lie algebra) using the Log-Euclidean metric (LEM):

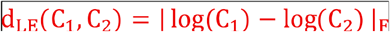

This metric preserves the fundamental structural relationships measured by AIRM but translates them into a mathematically flat coordinate system, preventing the ‘swelling effect’ and enabling exact vector calculus for trajectory tracking.

### Statistics

Unless stated otherwise, two-sided tests were used. Multiple testing was controlled by Benjamini– Hochberg FDR. For bar charts, bars represent mean ± SEM.

## Notes

### Competing Interest Statement

The authors have declared no competing interest.

